# Inferring the frequency spectrum of derived variants to quantify adaptive molecular evolution in protein-coding genes of *Drosophila melanogaster*

**DOI:** 10.1101/039404

**Authors:** Peter D. Keightley, José L. Campos, Tom R. Booker, Brian Charlesworth

**Affiliations:** Institute of Evolutionary Biology, School of Biological Sciences, University of Edinburgh, Edinburgh EH9 3FL, United Kingdom

## Abstract

Many approaches for inferring adaptive molecular evolution analyze the unfolded site frequency spectrum (SFS), a vector of counts of sites with different numbers of copies of derived alleles in a sample of alleles from a population. Accurate inference of the high copy number elements of the SFS is difficult, however, because of misassignment of alleles as derived *versus* ancestral. This is a known problem with parsimony using outgroup species. Here, we show that the problem is particularly serious if there is variation in the substitution rate among sites brought about by variation in selective constraint levels. We present a new method for inferring the SFS using one or two outgroups, which attempts to overcome the problem of misassignment. We show that two outgroups are required for accurate estimation of the SFS if there is substantial variation in selective constraints, which is expected to be the case for nonsynonymous sites of protein-coding genes. We apply the method to estimate unfolded SFSs for synonymous and nonsynonymous sites from Phase 2 of the Drosophila Population Genomics Project. We use the unfolded spectra to estimate the frequency and strength of advantageous and deleterious mutations, and estimate that ˜50% of amino acid substitutions are positively selected, but that less than 0.5% of new amino acid mutations are beneficial, with a scaled selection strength of *N_e_s ≈* 12.

## Introduction

Most protein sequences are strongly conserved between species, which suggests that the majority of amino acid-changing mutations are selectively removed from populations (Graur and Li 2000). The nature of the selective forces acting on the mutations that become fixed between species is central for a variety of questions in population genetics. These include understanding the maintenance of variation within species, the causes of variation in nucleotide diversity across the genome and the nature of evolutionary adaptation. Evidence for pervasive selection in the genome comes from observations of positive correlations between nucleotide diversity at putatively neutrally evolving sites and the rate of recombination (Begun and Aquadro 1992), and negative correlations between local genomic diversity and the presence of functional elements (such as protein-coding exons or conserved noncoding elements; Cai et al 2009; Hernandez et al 2011; Lohmueller et al 2011; Halligan et al 2013; Enard et al 2014; Deinum et al 2015). These correlations are likely to be caused by natural selection acting on functional sites in the genome reducing diversity at linked sites, but the precise nature of the selective forces involved is unresolved, since both selective sweeps due to positive selection and background selection caused by purifying selection can contribute to these patterns.

One approach to discriminate between the contributions of neutral, deleterious and advantageous substitutions to molecular evolution is based on the McDonald-Kreitman test (McDonald and Kreitman 1991), which compares within-species polymorphism to between-species divergence. Initially conceived as a test of departure from neutrality in a specific gene, it was subsequently adapted to estimate the proportion of substitutions driven to fixation by positive selection between-species for a class of sites in the genome (Fay et al. 2002; Smith and Eyre-Walker 2002). It does not, however, directly provide information on the rate of occurrence of advantageous mutations or on the magnitude of their selective effects. Furthermore, the approach is compromised if there has been a demographic change altering the fixation probability of selected alleles (either advantageous or disadvantageous), the signature of which is not captured by analysis of the polymorphism data (Eyre-Walker 2002).

Other ways of combining polymorphism and divergence data, or focusing on polymorphism data only to infer genome-wide selection may be more fruitful. Andolfatto (2007) and Macpherson et al. (2007) showed that there is a negative correlation between synonymous site polymorphism and nonsynonymous divergence in Drosophila, and used this information to estimate the strength of selection and frequency of adaptive protein evolution. Both studies concluded that there is widespread adaptive evolution, but estimates of the strength of selection and frequency of adaptive substitution depended on the size of the genomic window considered in the analysis. A related approach fits a population genetic model to mean reductions in diversity observed around nonsynonymous sites that have experienced a substitution between related species (Sattath et al 2011). This does not depend on assuming a specific window size. The best-fitting model suggests that there is substantial variation in the fitness effects of adaptive amino substitutions in Drosophila, potentially shedding light on the different results of Andolfatto (2007) and Macpherson et al. (2007).

We have previously described an approach that attempts to simultaneously infer the rate and strength of deleterious and beneficial mutations occurring in a class of sites in the genome, which exploits the shape of unfolded site frequency spectrum (uSFS) (Schneider et al 2011). The uSFS is a vector of counts of sites that have *j* copies of the derived allele, where 0 ≤ *j* ≤ *n*, and *n* is the number of copies in the sample. By using the uSFS, information for inferring the strength of selection comes mainly from current polymorphism within a focal species rather than divergence from an outgroup species. The first step is to infer demographic parameters using the SFS for quasi-neutrally evolving sites, such as synonymous sites. Conditioning on the estimates of the demographic parameters, selection parameters are estimated for a selected site class SFS (e.g., for nonsynonymous sites). These parameters describe the distribution of fitness effects (DFE) for deleterious mutations and the frequency of occurrence and strength of selection for one or more classes of advantageous mutations. Inferring adaptive evolution parameters requires that there is an excess of high frequency derived variants above and beyond that expected from demographic change and from negative selection acting on the bulk of mutations.

Applying the method of Schneider et al (2011), or any method that uses the frequencies of high frequency derived variants, therefore depends on accurate inference of the uSFS. Inference of the uSFS is potentially compromised, however, by misassignment of the ancestral state, and this tends to affect high frequency elements of the SFS disproportionately (Fay and Wu 2000; Baudry and Depaulis 2003; Hernandez et al 2007; De Maio et al. 2013; Glémin et al. 2015). Current methods to infer the uSFS rely on a single outgroup (Hernandez et al 2007) or require genome-wide polymorphism data from multiple species (De Maio et al. 2013). Schneider et al (2011) also described a method for inferring the uSFS, but we have recently determined that this tends to overestimate the frequency of high frequency derived variants (Halligan et al 2013). Here, we present a new method for inferring the uSFS, using information from one or two outgroup species that aims to address this problem, which we thoroughly test by simulations. We apply this method to a recent, whole-genome polymorphism data set for protein-coding genes from a sample of *Drosophila melanogaster* genomes originating from a Rwanda population close to their ancestral range. From the inferred uSFS, we estimate the frequencies and effects of deleterious and advantageous amino acid-changing mutations.

## Methods

### Inferring the uSFS - basic assumptions

A focal species is sequenced at multiple sites in a cohort of individuals sampled from a population. The possibility of more than two alleles segregating at a site in the focal species is disregarded. To infer the uSFS, we need to compute probabilities for the possible states of the allele ancestral to the observed alleles in the focal species (Figure 1a). We compute these probabilities using information from a single gene copy, assumed to be randomly sampled at each site, from either one or two outgroup species. Polymorphism in the common ancestor of the outgroup(s) and the focal species is disregarded; bias introduced by violating this assumption is investigated using simulations. Initially, we assume that all types of base substitution are equally frequent. Distinct transition and transversion rates are subsequently incorporated. The consequences of violating the equal mutation rates assumption in the basic method are explored by simulations.

**Figure 1:**
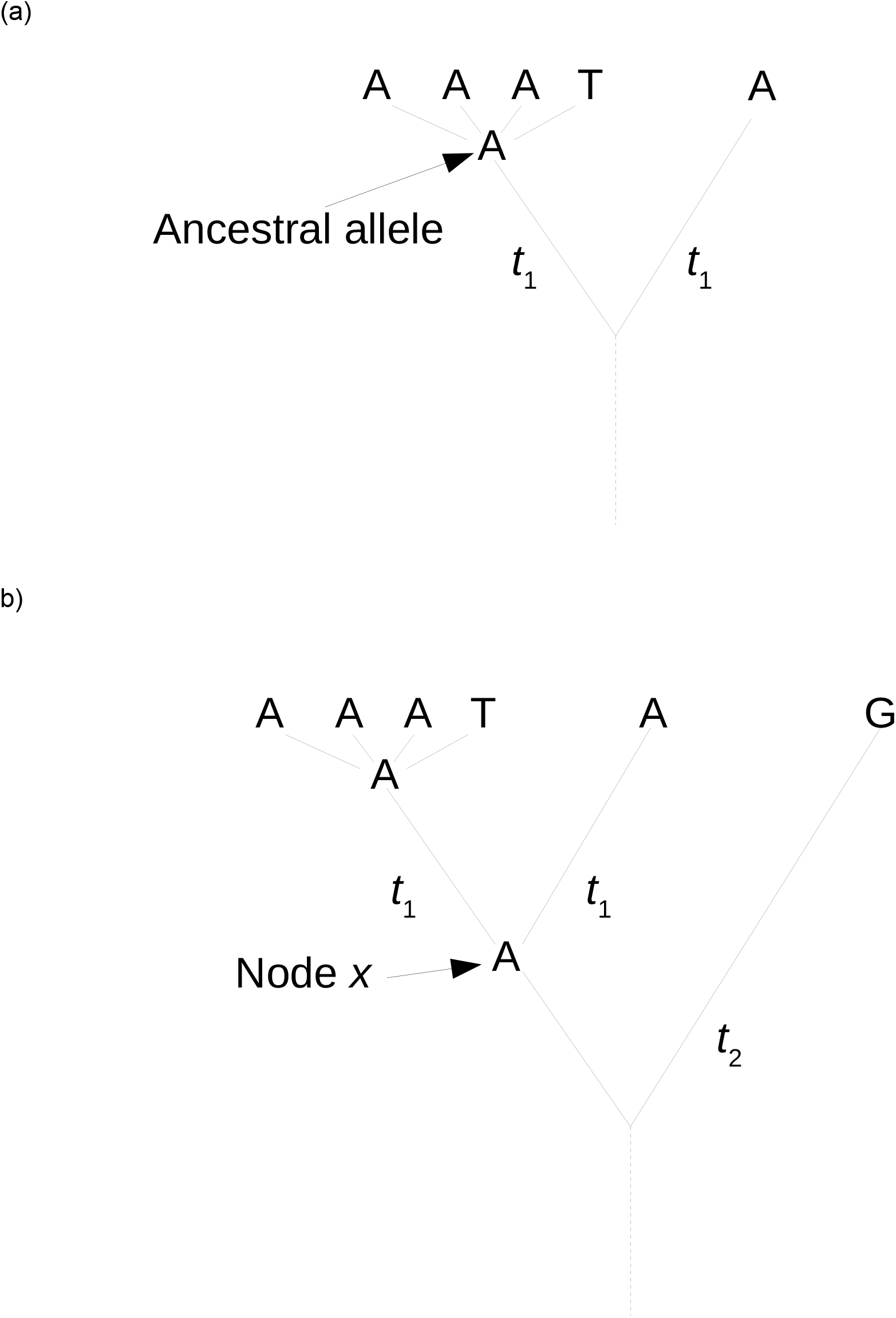
Example of a site at which four copies are sequenced in the focal species, where A is the major allele and the ancestral allele and T is the minor allele. (a) A single outgroup has the same state as the ancestral allele. (b) There are two outgroups and an internal node *x.* Time *t*_2_ is the total number of generations from *x* to outgroup 2.

### Single outgroup, single mutation rate parameter

Here, we illustrate the approach for inferring the uSFS assuming a single outgroup and a single evolutionary divergence parameter (*K*). This is the divergence between the allele ancestral to the observed allele(s) in the focal species and a single outgroup (Figure 1a). We do not need to consider mutations from the ancestral allele to the observed segregating alleles in the focal species. In this and the methods that follow (i.e., which allow different transition and transversion rates and two outgroups), a two-stage approach is implemented. First, the evolutionary divergence parameter(s) is estimated by maximum likelihood (ML). Second, assuming perfect knowledge of divergence parameter(s), the elements of the uSFS are estimated one-by-one by ML.

### Single outgroup stage 1 - ML estimation of K

Assume that the data consist of counts of numbers of different alleles observed in a sample of *n* copies in the focal species (n = 4 gene copies in Figure 1 and Table 1, for example) and a single copy from an outgroup species. Let *K* be the expected number of mutations distinguishing the allele ancestral to these 4 gene copies and the outgroup. Defining *x,* as the allelic configuration observed at site *i* in the focal species and the outgroup, and assuming independence among sites, the likelihood of the data for all sites combined is:

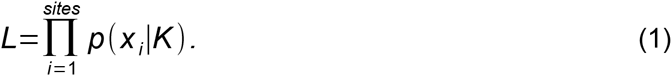

**Table 1.** Five possible configurations (*y*_1_… *y*_5_) of numbers of copies of alleles at a site observed in the focal species and the outgroup for the case of four copies sampled in the focal species. There is either no or one copy of a minor allele present in the focal species (T in this case). Assuming that there are from *m* = 0 to 2 mutations between the ancestral allele of the alleles present in the focal species and the outgroup (Figure 1), the conditional probabilities 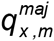 and 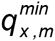 of observing configuration *x*, given that the ancestral allele is the major or the minor allele, respectively, are shown.

| **Observed state** |  |  |  | **Conditional probability** |  |  |
| --- | --- | --- | --- | --- | --- | --- |
| **Configuration** | **Focal species** | **Outgroup** | **$m = \text{no. mutations}$** | $q_{x,m}^{maj}$ | $q_{x,m}^{min}$ | $q_{x,m} = q_{x,m}^{maj} + q_{x,m}^{min}$ |
| $y_1$ | AAAA | A | 0 | 1 | 0 | 1 |
|  |  |  | 1 | 0 | 0 | 0 |
|  |  |  | 2 | 1/3 | 0 | 1/3 |
| $y_2$ | AAAA | T | 0 | 0 | 0 | 0 |
|  |  |  | 1 | 0 | 1 | 1 |
|  |  |  | 2 | 0 | 2/3 | 2/3 |
| $y_3$ | AAAT | A | 0 | 1 | 0 | 1 |
|  |  |  | 1 | 0 | 1/3 | 1/3 |
|  |  |  | 2 | 1/3 | 2/9 | 5/9 |
| $y_4$ | AAAT | T | 0 | 0 | 1 | 1 |
|  |  |  | 1 | 1/3 | 0 | 1/3 |
|  |  |  | 2 | 2/9 | 1/3 | 5/9 |
| $y_5$ | AAAT | C | 0 | 0 | 0 | 0 |
|  |  |  | 1 | 2/3 | 2/3 | 4/3 |
|  |  |  | 2 | 4/9 | 4/9 | 8/9 |

If the focal species is monomorphic there are two possible configurations of alleles (*y*_1_ and *y*_2_; Table 1) and there are three configurations if the focal species is polymorphic (*y*_3_, *y*_4_ and *y*_5_; Table 1). Noting the symmetry of configurations *y*_3_ and *y*_4_, equation (1) can therefore be rewritten as:

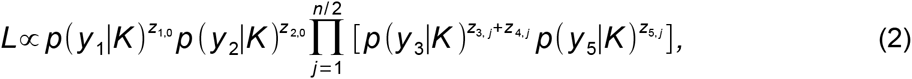

where *Z_xj_* is the number of sites showing the configuration with subscript *x*, given that there are *j* minor allele copies in the focal species.

Assuming that the number of mutations (*m*) is Poisson distributed with probability *P*(*m|K*), the probability of configuration *y_x_*, given divergence *K* is:

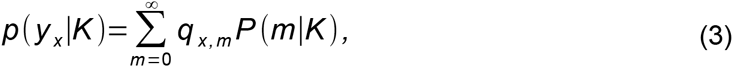

where *q_x,m_* is the conditional probability for allelic configuration *y_x_*, given that there are *m* mutations (Table 1). In practice, we considered only up to two mutations in the summation in equation (3). Simulations with *K* up to 20% suggested that allowing more that two mutations had a negligible effect on estimates of *K* or the SFS elements.

For example, the probability (*q*_1,0_) of observing configuration *y*_1_ if there have been no mutations is equal to 1; it is not possible to observe configuration *y*_1_ if there has been one mutation (i.e., *q*_1,1_ = 0); *q*_1,2_ = 1/3 because if there had been two mutations, nucleotide A could have mutated to any other nucleotide and then must have mutated back to A. The natural log likelihood with respect to *K*, i.e., log(equation 2), was maximized by the Golden Search algorithm (Press et al. 1992).

### Single outgroup stage 2 - ML estimation of the uSFS elements, given K

The approach is to find the ML estimate of the proportion of probability density, *π_j_*, attributable to the major allele being ancestral *versus* the minor allele being ancestral for each element of the SFS, while assuming the fixed ML estimate of *K* from stage 1. There are therefore *n*/2 + 1 ML estimates to be made. To compute the likelihood of *π_j_*, we need to consider sites for which there are *j* copies of one allele and *n* − *j* copies of a different allele in the sample of *n* copies. For invariant sites (*j* = 0), there are *Z*_1,0_ and *Z*_2,0_ sites that have allelic configurations *y*_1_ and*y*_2_, respectively (Table 1). For variant sites (*j* ≠ 0), there are three possible allelic configurations (*y*_3_, *y*_4_ and *y*_5_; Table 1), but sites where the outgroup allele is different from the copies observed in the focal species (configuration *y*_5_) provide no information about the uSFS, and so can be disregarded. Note that these sites *do* contribute to the estimate of *K*. We therefore have *Z*_3*j*_ and *Z*_4*j*_ sites with the two informative configurations. The likelihood for variant sites that have *j* minor alleles in the focal species is:

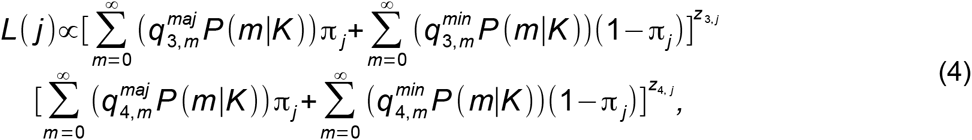

where superscript *maj* (*min*) on *q* implies that the ancestral allele is the major (minor) allele (Table 1). The likelihood for invariant sites in the focal species is:

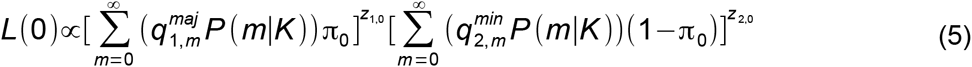

We considered only up to two mutations in the summations in (4) and (5). Log likelihood with respect to each *π_j_* was maximized by the Golden Search algorithm.

This method can be adapted to infer uSFS elements where there are different transition and transversion rates estimated (see Supplementary Information; Table S1), and where there are two outgroups (see Supplementary Information; Table S2).

### Simulations

We assessed the performance of the uSFS inference procedures using Monte Carlo simulations incorporating ancestral polymorphism and unequal transition/transversion rates. We analysed data sets containing large numbers of sites specifying the allelic states for *n* copies sampled from the population of a focal species and single copies sampled from populations of one or two outgroup species. The simulated populations were diploid and of constant population size *N* = 100. We generally assumed that the neutral diversity *θ* is equal to 0.01 by setting the mutation rate per site per generation to *μ* = θ/4*N*. We simulated unlinked nucleotide sites that could be in of one four states (A, T, C, G). An ancestral population was initiated with equal frequencies of the four nucleotides, and allowed to evolve to mutation-drift equilibrium for 20N burn-in generations. A site of an individual was mutated with probability *μ* each generation by randomly altering its current nucleotide state. In general, the probability of a transition mutation was equal to 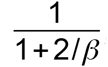, where *β* > 1 implies there is transition:transversion mutation bias. In the case of a single outgroup, two separate populations were evolved for *t*_1_ generations after the burn-in to produce a focal population and an outgroup population (Fig 1a). When simulating two outgroups, an outgroup 2 population was evolved for (*t*_1_ + *t*_2_)/2 generations and a second population was evolved for (*t*_2_ − *t*_1_)/2 generations up to node *x* (see Fig 1b). Two populations were then evolved from the node *x* population for *t*_1_ generations to produce a focal population and an outgroup 1 population.

In many simulations we assumed that all sites evolve neutrally. We also simulated variation in the rate of substitution among sites caused by variation in the strength of purifying selection. A fraction *C* of sites was designated as selectively constrained sites. Any allele different in state from the wild type allele that arose at such sites was designated as mutant and had a selective disadvantage *s*/2. Effects on fitness were multiplicative. Fertility selection was carried out by sampling individuals for reproduction with replacement in proportion to their relative fitness.

We quantified bias (in %) affecting estimates of elements of the SFS as the percentage deviation from the true value of that element. We also estimated the scaled root mean squared error (RMSE in %) for elements of the SFS. RMSE incorporates variance among estimates (since one method might produce less variable estimates of SFS elements about the true values than another) and is also influenced by bias:

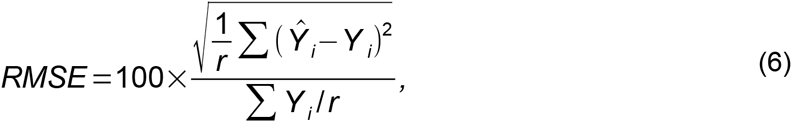

where *Ŷ_i_* is the estimate for an SFS element from simulation replicate *i* and *Y_i_* is the corresponding true value of that element, and *r* is the number of simulation replicates.

### D. melanogaster potymorphism data

We obtained polymorphism data from an African population of *Drosophila melanogaster* comprising 17 Rwandan haploid genomes (RG18N, RG19, RG2, RG22, RG24, RG25, RG28, RG3, RG32N, RG33, RG34, RG36, RG38N, RG4N, RG5, RG7 and RG9), which have been estimated to have the lowest levels of admixture with European populations (less than 3%, see Figure 3b of Pool et al., 2012). We downloaded FASTQ files from the *Drosophila* Population Genomics Project (http://www.dpgp.org/dpgp2/candidate/). We further masked any regions of the African samples with evidence of admixture from European populations, using the admixture coordinates reported by Pool et al. (2012). Following Pool et al. (2012), sites with a BWA quality score below *Q* = 31 (equivalent to a PHRED score of 48, and approximately equivalent to one error per 100 kb) were also masked. This produced the Q31 data set, which is the focus of most of the analysis. We also analysed a more stringently filtered Q41 data set. From the FASTQ files, we extracted protein-coding regions, using gene annotations from FlyBase release version 5.33 (www.flybase.org) and made FASTA files containing all samples (17 copies), we excluded genes within non-crossing over regions (see Campos et al. 2012). For each *D. melanogaster* gene with multiple transcripts, we chose one transcript at random. We included as outgroups the orthologous genes of *D. simulans* (r2.01) and *D. tyakuba* (r1.3), obtained from the *D. melanogaster-D. simulans-D. tyakuba* gene alignments of Hu et al. (2013), from which we selected the coding regions corresponding to our selected transcripts.

### Estimating the DFE and the rate and strength of adaptive mutations along with the frequencty of adaptive substitutions

Using the inferred unfolded uSFSs for two outgroups and with no transition/transversion bias assumed, we estimated parameters of the DFE and adaptive mutations by the ML method described by Schneider et al (2011), which is incorporated into the software DFE-alpha (Keightley and Eyre-Walker 2009), with the following modifications. We first fitted a three-epoch demographic model to the neutral (i.e., synonymous) uSFS, allowing two changes of population size, first from *N*_1_ to *N*_2_, then from *N*_2_ to *N*_3_ at times *t*_2_ and *t*_3_, respectively, while also fitting parameters specifying the fractions of unmutated sites (*f*_0_) and sites fixed by drift (*f*_2*N*_). By fitting this model to the DPGP synonymous uSFS, we found that high frequency elements were under-predicted. The estimated uSFS for synonymous sites contains a small uplift in the last element, which cannot be explained under the demographic and mutational model fitted. This uplift could reflect hitchhiking with selected amino acid variants or positive selection on synonymous variants. Alternatively, it could be caused by residual misassignment of low frequency variants. We assumed that such processes also affected the nonsynonymous uSFS, and would lead to upwardly biased estimates of positive selection parameters if not corrected. In a similar manner to that described by Glémin et al (2015), which follows the approach of Eyre-Walker et al (2006), we therefore corrected elements of the nonsynonymous uSFS (*N_j_*) using the deviations of the observed (*S_j_*) from the fitted (*E_j_*) elements of the synonymous uSFS:

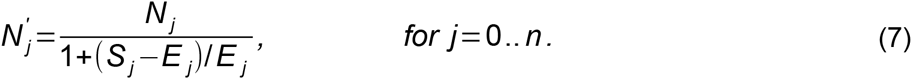

We assessed goodness of fit by comparing fitted uSFSs to observed uSFSs using a χ^2^ statistic, but because the numbers of sites in derived class *j* and ancestral class *n − j* are non-independent, we do not perform formal significance tests.

Conditioning on the values of the parameters fitted to the synonymous SFS, parameters specifying the effects and relative frequencies of deleterious and advantageous mutations were fitted by ML to the corrected nonsynonymous uSFS. We either assumed that the fitness effects of deleterious mutations were drawn from a gamma distribution (which is specified by a shape and a scale parameter) or, following Kousathanas and Keightley (2013), that there were *n_d_* fixed classes of deleterious mutations, where the fitness effect and frequency of class *i* are *s_d,i_*. and *p_d,i_* respectively. We fitted *n_a_* classes of advantageous mutations, where the fitness effect and frequency of class *J* are *s_a,j_*. and *p_a,j_*, respectively, such that 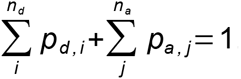. The gamma DFE represents a single, continuously variable class of deleterious mutations.

To find maximum likelihood estimates (MLEs) we carried out runs with large numbers of combinations of random starting values. We estimated 95% confidence limits for the proportion of adaptive mutations and their selective strength from profile likelihoods on the basis of drops in log likelihood of 2 units from their respective MLs. For each point in each profile likelihood we used the highest likelihood obtained from 20 runs using different starting values sampled around the MLEs. Estimates of α, the proportion of adaptive substitutions, and *ω_a_,* the rate of adaptive substitution relative to the rate of neutral substitution, were obtained as described by Schneider et al (2011).

## Results

### Simulations - single outgroup

To investigate the performance of the uSFS inference procedure under circumstances where the data closely conform to the assumptions of the model, we simulated a focal population and a single outgroup with nucleotide divergence *K* = 0.1, no transition/transversion bias (*β* = 1) and no selection. We assumed that *θ* = 4*N_e_μ* = 0.01, so that *θ* << *K* and few polymorphic sites in the focal species are also polymorphic in the ancestral population prior to the split between the focal species and the outgroup. Figure 2 shows the true uSFS (calculated using knowledge of the ancestral state for each site) and the uSFSs inferred using the single outgroup method described here and the method of Schneider et al. (2011). The new approach is therefore capable of estimating the uSFS with little bias on average, including high frequency elements of the SFS. The method of Schneider et al. (2011) tends to overestimate high frequency SFS elements, presumably because polymorphic sites having an outgroup allele inconsistent with either allele present in the focal species are mis-assigned. Our new approach appears to give nearly unbiased estimates of the uSFS elements as long as the divergence to the outgroup is *K* < 0.3 (Figure S1).

**Figure 2:**
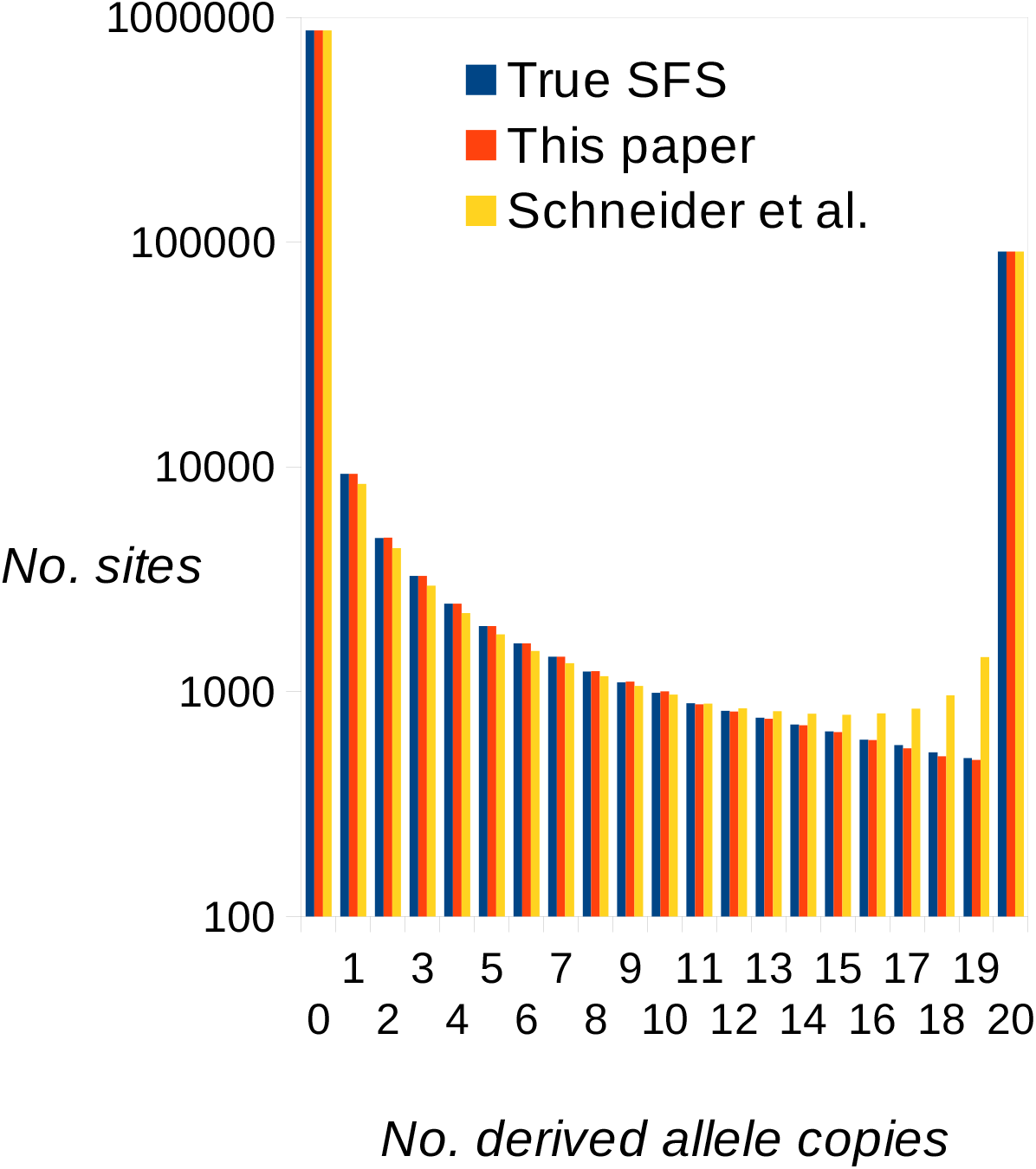
True uSFS (from simulation) and estimated uSFSs computed by the present method and by the method of Schneider et al. (2011), both using a single outgroup. 20 copies were sampled at each site of the focal species. Diversity *θ* = 4*Nμ* = 0.01, and divergence between the focal species and the outgroup was *K* = 0.1. There were 8 replicate simulations, each with 10^6^ sites, resulting in a negligible sampling variance for elements of the estimated uSFSs.

We extended the method to include the estimation of separate transition and transversion rate parameters (Supplementary Information; Table S1). This was tested by simulations of neutrally evolving sites, and also produces close to unbiased estimates of the uSFS in the presence of transition:transversion mutational bias (Figure S2a). The single rate parameter method also produces reasonably unbiased estimates of the uSFS unless there is substantial bias towards transition mutations, or the divergence is 0.2 or greater (Figure S2b).

### Simulations - two outgroups, neutrally evolving sites

We then compared the performances of the uSFS inference procedures allowing one or two outgroups. The results suggest that there is a clear benefit from using a second outgroup in terms of lower variance among replicates (lower RMSE) (Figure 3), but potentially a cost in terms of higher bias (i.e., there is a tendency for underestimation of the high frequency SFS elements), especially if the divergence from the second outgroup is small (Figure 3).

**Figure 3:**
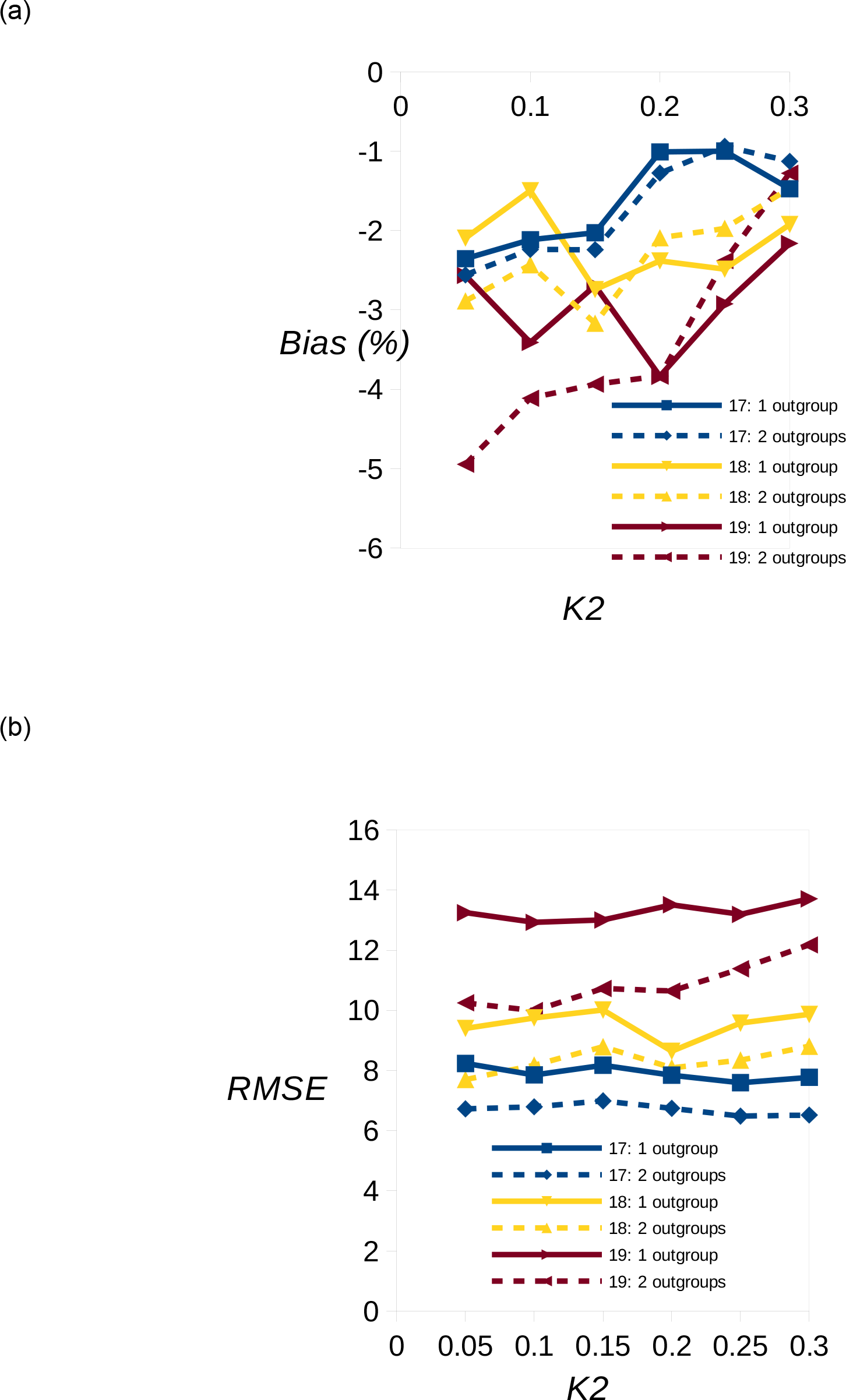
Estimated bias (%) (a) and RMSE (b) for estimates of uSFS elements 17, 18 and 19 plotted against divergence (*K*_2_) between node x and outgroup 2 (see Figure 1) for the case of 20 copies sampled at each site of the focal species. The solid and dotted lines show inferences using one or two outgroups, respectively. The divergence between the focal species and outgroup 1 was *K*_1_ = 0.1, and diversity in the focal species was θ = 0.01. There was no transition:transversion bias. 10^5^ sites were simulated in each of 160 replicates.

### Simulations - variable strength of purifying selection among sites

We then investigated the performance of the uSFS inference procedures in the presence of variation in the substitution rate and diversity among sites caused by purifying selection. We simulated this variation by assuming that a fraction *C* of sites are subject to negative selection (with selective disadvantage *s* = 0.1; see Materials and Methods), the remainder evolving neutrally. We found that with *C* ≈ 0.85 and *Ns* = 10 (so that mutant alleles rarely become fixed), divergence, diversity and the shape of the SFS simulated are similar to what we observe in the *D. melanogaster* polymorphism data for nonsynonymous sites, although in this case we assume a constant population size.

We compared the accuracy of the inferred uSFS using one or two outgroups, focusing on the high copy number elements of the SFS, which are hardest to estimate accurately. We assumed a neutral divergence between the focal species and the first outgroup of *K*_1_ = 0.1 (which is similar to the *D. melanogaster - simulans* divergence) and a neutral divergence between the internal node and the second outgroup of *K*_2_ = 0.15. The results suggest that there is a clear benefit, both in terms of reduced bias and reduced RMSE from using the information from a second outgroup (Figure 4). We see a only a small amount bias and reduced RMSE for a range of *C* values. Using information from a single outgroup, however, can lead to serious over-estimation of the high copy number SFS elements (as much as 15% in the cases shown). Presumably, the benefit of using a second outgroup applies when there are other sources of variation in the substitution rate among sites, such as variation in the mutation rate.

**Figure 4:**
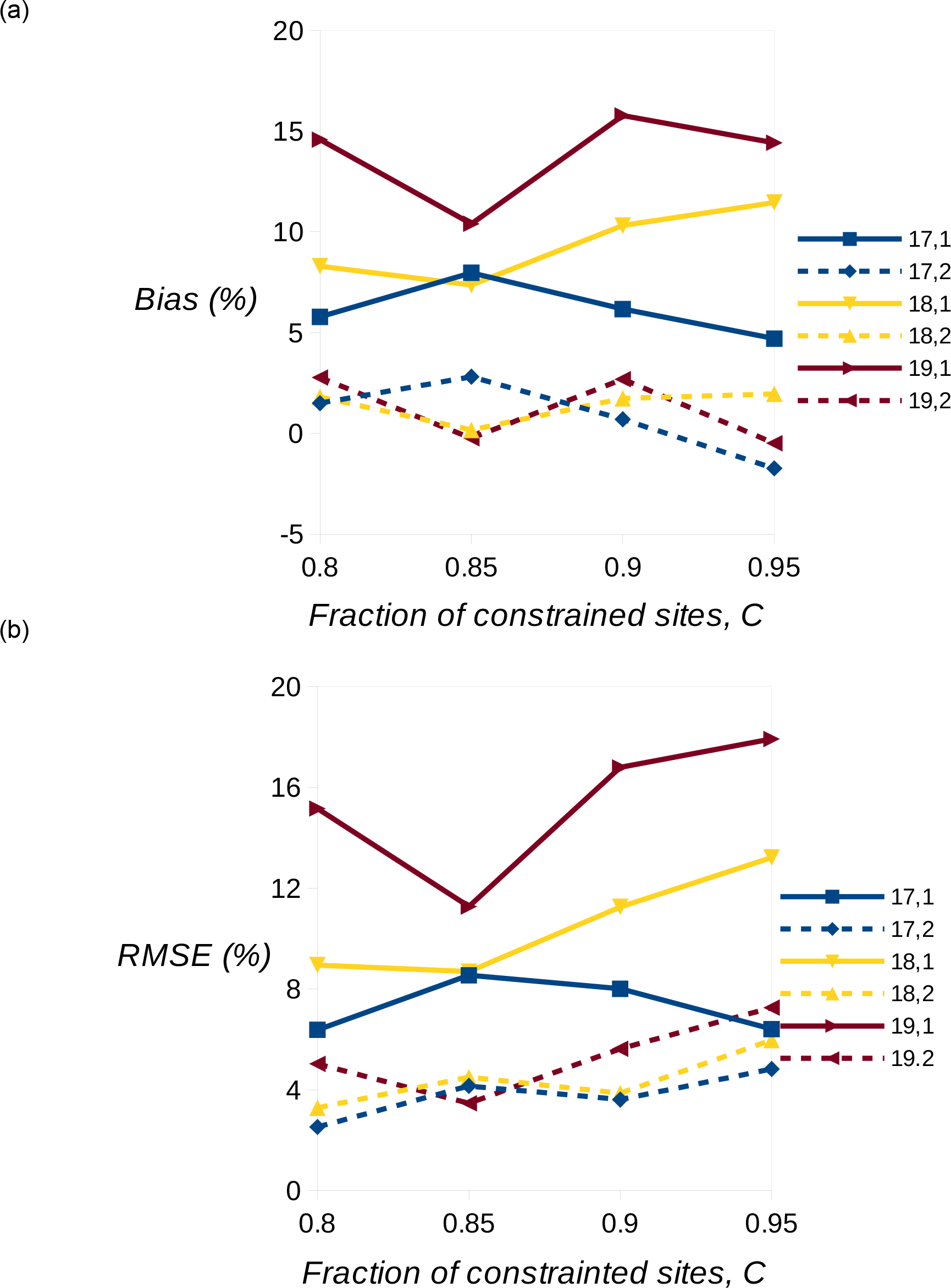
One versus two outgroup comparison in the presence of variation among sites in selective constraints. The panels show bias (a) and RMSE (b) in the last three elements of the uSFS. For example, the label 17,1 refers to the bias/RMSE affecting element 17 with one outgroup. A fraction *C* of sites were simulated with scaled selection coefficient *Ns* = 10 and the remainder evolve neutrally. 10^5^ sites were simulated per replication and 240 replicates. The divergence parameters for neutral alleles were *K*_1_ = 0.1 and *K*_2_ = 0.15.

### Inference of uSFSs and frequency and strength of adaptive molecular evolution in the D. melanogaster proteome

We applied the uSFS inference procedure to the polymorphism data set of protein-coding genes of the *D. melanogaster* DPGP Phase 2. Using two outgroups (D. *simulans* and *D. yakuba*), we inferred uSFSs for 4-fold and 0-fold sites (Figure 5). As expected, nucleotide diversity at 0-fold sites is substantially lower than that at 4-fold sites, and there is an enrichment of 0-fold singletons, consistent with negative selection acting on many nonsynonymous sites.

**Figure 5:**
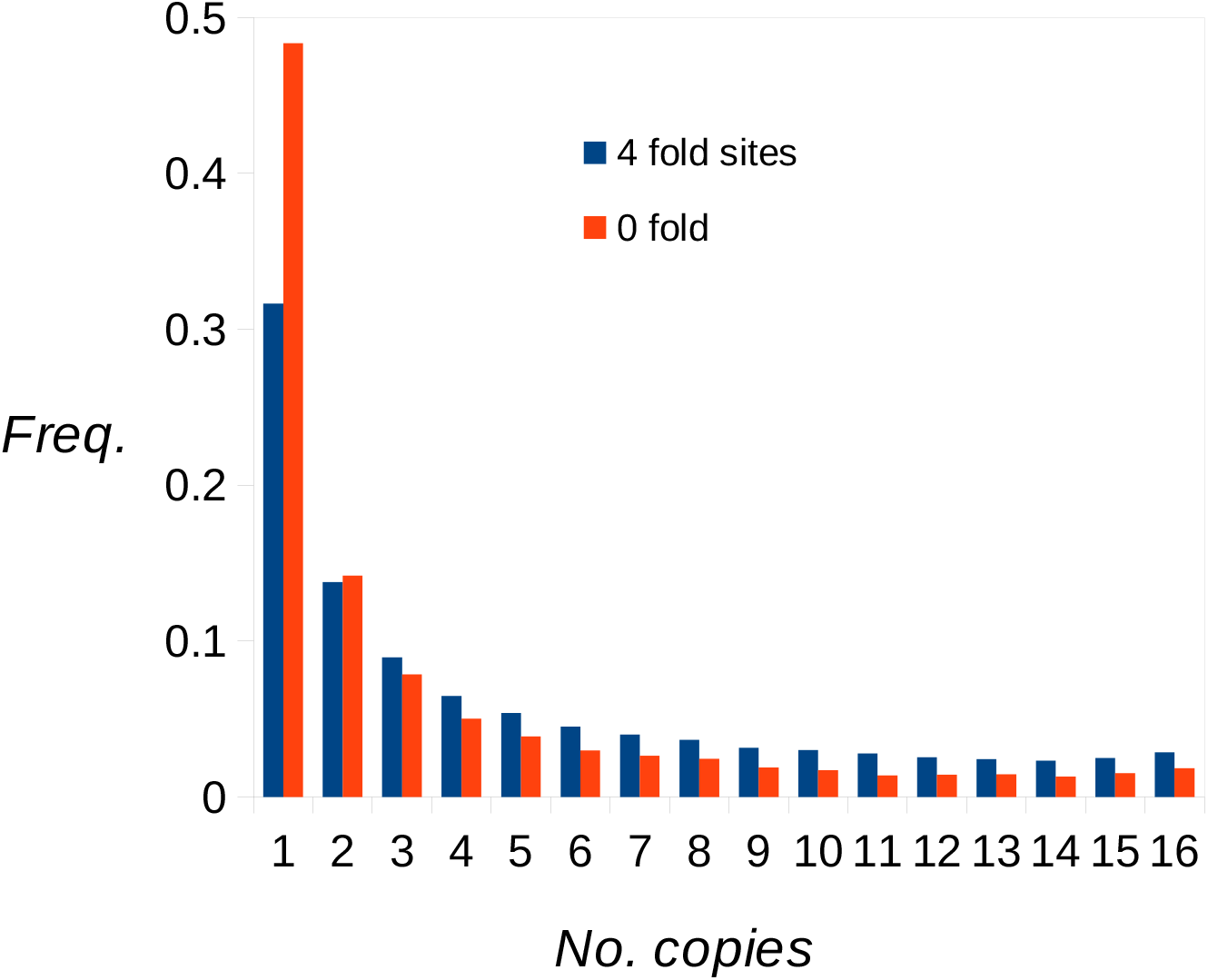
Unfolded SFSs for 0-fold and 4-fold sites of *D. melanogaster* protein-coding genes inferred using two outgroups (*D. simulans and D. yakuba*).

We then applied the approach of Schneider et al (2011) to estimate the rate of occurrence and selective strength of adaptive amino acid mutations. We fitted parameters of a three-epoch demographic model to the synonymous site data (Table S3; Figure S3); this model fitted much better than a two-epoch model (log likelihood difference = 221), and suggests that there was a population size bottleneck followed by a population expansion. There is, however, an appreciable deviation between the observed and fitted synonymous uSFS, particularly affecting the last element (Figure S3; χ^2^(16) = 138). We assumed that misinference would also affect the nonsynonymous uSFS, potentially leading to spurious estimates of adaptive molecular evolution. We therefore corrected the nonsynonymous uSFS using the deviation between the observed and fitted synonymous uSFS, as described in Methods. Uncorrected and corrected nonsynonymous uSFSs are shown in Figure S4.

Given the demographic parameter estimates from the synonymous site data, we then estimated parameters of the distribution of fitness effects (DFE) for deleterious mutations and the proportion (*p_a_*) and scaled selection strength acting on one or more classes of adaptive mutations (*N_e_s_a_*). Several models had similar levels of statistical support (Table 2). The best-fitting model gives an excellent fit to the data (Figure S4; χ^2^(16) = 16.9), and consisted of four classes of mutational effects: two classes of deleterious mutations, a class of neutral mutations, and a single class of advantageous mutations. There is substantial support for models with adaptive mutations (ΔsogL = 93 between the best fitting model and the same model excluding adaptive mutations). Assuming the four-class model, ML estimates of the proportion of advantageous mutations and the scaled strength of selection acting on them are *p_a_* = 0.0045 (approx. upper 95% CI = 0.012) and *Nesa* = 11.5 (approx. lower 95% CI = 5), respectively. Note that *p_a_* and *s_a_* are hard to estimate separately, but their product is well estimated. Other models that explain the data almost as well (gamma DFE, three classes of mutational effects) give somewhat different ML estimates of *p_a_* and *N_e_s_a_,* but the products of *p_a_* x *N_e_s_a_* are of similar magnitude (Table 2). Fitting additional classes of mutations (advantageous or deleterious) did not lead to a further increase in log likelihood. We then estimated the frequency of adaptive substitutions (*α*) and the rate of adaptive substitution relative to that of neutral substitution (*ω_a_*) from the proportions and fixation probabilities of the advantageous, neutral and deleterious mutation classes. The estimates are *α* = 0.57 and *ω_a_* = 0.096 for the 4-class model, but these are sensitive to the model assumed (Table 2).

**Table 2.** ML estimates of parameters from DFE-alpha for three different models involving different numbers of classes of mutational effects and a gamma DFE and the change in log likelihood (*Δ/ogL*) from the best-fitting model.

| *Parameter* | *ML estimates* |  |  |
| --- | --- | --- | --- |
|  | *4 class* | *3 class* | *Gamma* |
| $\beta$ | - | - | 0.35 |
| $p_{d1}$ | 0.88 | 0.89 | 1 |
| $N_e s_{d1}$ | -177 | -167 | -2120 |
| $p_{d2}$ | 0.076 | 0.10 | - |
| $N_e s_{d2}$ | -2.8 | -1.4 | - |
| $p_{d3}$ | 0.039 | - | - |
| $N_e s_{d3}$ | 0 | - | - |
| $p_a$ | 0.0045 | 0.0093 | 0.0031 |
| $N_e s_a$ | 11.5 | 6.7 | 17 |
| $\alpha$ | 0.57 | 0.89 | 0.68 |
| $\omega_a$ | 0.096 | 0.14 | 0.091 |
| $\Delta\log L$ | 0 | -0.6 | -2.1 |

## Discussion

There were three main motivations for this study. First, we had determined that a previously described method to infer the uSFS (Schneider et al 2011) tends to overestimate high frequency SFS elements. Second, using parsimony for inferring ancestral states of high frequency elements of the SFS is problematical, because the corresponding low frequency elements usually involve a far greater number of sites, and these tend to be misassigned as high frequency elements, potentially leading to an over-estimation of the frequency of alleles under positive selection. Third, large, genome-wide polymorphism data sets offer the opportunity to investigate the frequency and strength of ongoing adaptive molecular evolution, using a method also described by Schneider et al (2011), but this requires accurate inference of the uSFS. The development of this approach was motivated by inconsistent results emerging from the application of variants of the McDonald-Kreitman (MK) test, such as the methods of Welch (2006), DFE-alpha (Eyre-Walker and Keightley 2009) and DoFE (built on Eyre-Walker et al 2006).

These methods all estimate the frequency of adaptive substitutions in a set of loci by contrasting polymorphism data in a focal species with divergence from an outgroup species. For example, many estimates of the proportion of adaptive amino acid substitutions (a) in plants are negative, some significantly, at face value implying that there is little adaptive protein evolution (Gossmann et al 2010). The true value of *α* cannot be negative, however, and this result may reflect the presence of widespread population structure in plant species, which distorts the SFS and could bias estimates of *α* downwards. Some estimates of *α* in great apes are also negative (Good et al 2013). A clear example of inconsistency comes from a reciprocal analysis of genome-wide polymorphism data in murid rodents, where an estimate of *α* in wild house mice using divergence from the rat is strongly and significantly positive, i.e., *α* ≈ 0.3 (Halligan et al 2013), whereas an estimate using polymorphism within wild brown rats and divergence from the mouse is strongly and significantly negative, i.e., *α* ≈ -0.3 (Deinum et al 2015). The negative estimate presumably reflects a recent population bottleneck in the brown rat, leading to over-prediction of the number of fixed slightly deleterious mutations.

In contrast to McDonald-Kreitman-based methods, the method of Schneider et al (2011) uses information on polymorphism data within a species to infer ongoing adaptive molecular evolution. It can be set up to use no information from sites fixed for the derived allele, but we did not do that here. By simulations, we investigated the circumstances under which accurate inference of the uSFS is possible. The most important potential source of misinference we identified is variation in the substitution rate, affecting the joint spectrum of polymorphism in the focal species and divergence(s) from the outgroup(s). This could either be due to variation in the mutation rate between different kinds of sites or variation between sites in selective constraints or adaptive potential. In principle, it is possible to account for some components of variation in the mutation rate by explicit modeling (e.g., transition-transversion bias). Selection that varies among sites appears to be a more important issue, however, and is more difficult to model. Our simulations show that with a single outgroup only, high copy number uSFS elements are potentially seriously over-estimated if there is variation in selective constraints among sites. This is because the divergence between the ancestral allele and the outgroup is computed as an average across sites, but this will be lower than the divergence at the subset of unconstrained sites, so multiple hits are under-corrected at these sites. Our simulation results suggest that incorporating a second outgroup substantially corrects this problem, allowing accurate estimation of the uSFS. It should be feasible to extend our approach to include multiple outgroups, although there are presumably diminishing returns and potential biases from adding more distant outgroups.

We applied our new uSFS inference approach to the *D. melanogaster* DPGP Phase 2 data for protein-coding genes, and several aspects of the results are noteworthy. The inferred uSFS for synonymous sites contains a small, but appreciable uplift in the last element (Figure S3). With the demographic and mutational models fitted by DFE-alpha, however, it is not possible to obtain an uplift in the fitted uSFS. The apparent increase in the frequency of high copy number derived synonymous alleles could be genuine, and explained, for example, by hitchhiking with selected amino acid variants or selection on synonymous variants (Zeng 2010; Clemente and Vogl, 2012; Lawrie et al 2013). Alternatively, it could be caused by residual misassignment of low frequency variants. We investigated whether this might be due to sequencing errors by analyzing a more stringent set of SNPs (Q41). The inferred uSFSs are extremely similar to the uSFSs analysed (Q31; Figure S5), suggesting that sequencing errors in DPGP are not an important source of misinference. We corrected the nonsynonymous uSFS based on the deviation from the fitted and observed synonymous uSFS.

Fitting the demographic parameters estimated from synonymous sites, then estimating selection parameters by ML, resulted in a close fit to the corrected nonsynonymous uSFS (Figure S4), but several alternative models also give excellent fits (Table 2). Taking the best-fitting model at face value, the results therefore imply that there is a major contribution from adaptive amino acid substitutions to protein evolution in *D. melanogaster,* i.e., *α* ≈ 0.5. This figure is consistent with several studies employing variants of the McDonald-Kreitman test to estimate the frequency of adaptive protein evolution (Fay et al 2002; Smith and Eyre-Walker 2002; Welch 2006; Andolfatto 2007; Eyre-Walker and Keightley 2009; Campos et al 2014). The estimated selective effects of adaptive mutations are also consistent with estimates for the more common, weakly selected of the two classes inferred by Sattath et al (2011), based on changes in diversity around substituted nonsynonymous sites. However, Sattath et al estimated that only about 13% of amino acid substitutions cause selective sweeps, arguing that this low value could reflect a prevalence of partial sweeps. On the other hand, Schneider et al (2011) used information from high frequency polymorphisms, which is most relevant for inferring the ongoing strength of selection and the frequency of relatively weakly selected variants. This is because strongly selected mutations are expected to be relatively rare and sweep rapidly to fixation, leaving little detectable footprint in the uSFS.

